# State-dependent role of interhemispheric pathway for motor recovery in primates

**DOI:** 10.1101/2023.04.23.538021

**Authors:** Masahiro Mitsuhashi, Reona Yamaguchi, Toshinari Kawasaki, Satoko Ueno, Yiping Sun, Kaoru Isa, Jun Takahashi, Kenta Kobayashi, Hirotaka Onoe, Ryosuke Takahashi, Tadashi Isa

**Affiliations:** Department of Neuroscience, Graduate School of Medicine, Kyoto University, Kyoto, 606-8501, Japan; Department of Neurology, Graduate School of Medicine, Kyoto University, Kyoto, 606-8507, Japan; Institute for the Advanced Study of Human Biology (WPI-ASHBi), Kyoto University, Kyoto, 606-8501, Japan; Department of Neurosurgery, Graduate School of Medicine, Kyoto University, Kyoto, 606-8507, Japan; Department of Clinical Application, Center for iPS Cell Research and Application, Kyoto University, Kyoto 606-8507, Japan; Section of Viral Vector Development, National Institute for Physiological Sciences, Okazaki, 444-8585, Japan; Graduate University of Advanced Studies (SOKENDAI), Hayama, 240-0193, Japan; Human Brain Research Center, Graduate School of Medicine, Kyoto University, Kyoto 606-8397, Japan

## Abstract

Whether and how the interhemispheric pathway is involved in post-injury motor recovery is controversial. Unidirectional chemogenetic blockade of the interhemispheric pathway from the contralesional to ipsilesional premotor cortex impaired dexterous hand movements during the early recovery stage after lesioning the lateral corticospinal tract in macaques. Furthermore, electrocorticographical recording showed that the low frequency band activity of the ipsilesional premotor cortex around the movement onset was decreased by the blockade during the early recovery stage, while it was increased by blockade during the intact state and the late recovery stage. These results demonstrate that the action of the interhemispheric pathway changed from inhibition to facilitation, leading to the involvement of the ipsilesional sensorimotor cortex in hand movements during the early recovery stage. The present study offers new insights into the state-dependent role of the interhemispheric pathway and a therapeutic target in the early recovery stage after lesioning of the corticospinal tract.

---

The lateral corticospinal tract (l-CST) is crucial for dexterous hand movements in higher primates^1,2^. Lesions of the l-CST impair hand dexterity and the activities of daily living. It is widely known that damaged neurons regenerate rarely in adults^3,4^, but impaired motor function can be recovered considerably by rehabilitative training^5,6^. Thus, the mechanism of functional recovery is associated with the training-induced plasticity of residual neural circuits^7,8^. Identifying the pathways for functional recovery would reveal targets for future neuromodulation therapies.

We previously demonstrated that after l-CST lesioning of the middle cervical cord (C4–C5), the ipsilesional sensorimotor cortex is activated during the early recovery stage by using positron emission tomography, and that inactivation of the ipsilesional primary motor cortex (M1) with muscimol affects the recovery of hand dexterity^9^. Thus, the contribution of the ipsilesional motor cortex to the early recovery stage is clear; however, which pathway activates the ipsilesional sensorimotor cortex remains elusive. Then, Chao et al. performed Granger causality (GC) analysis, which can provide an unidirectional causal dependence between two signals, and evaluated the connectivity between all pairs of multichannel (30 × 30 channels) electrocorticography (ECoG) records spanning the bilateral sensorimotor cortices that were obtained throughout the intact and recovery periods using a multidimensional algorithm for dimension reduction^10^. Their results suggested that interhemispheric signal flow from the contralesional to ipsilesional premotor cortex (PM) at the α and low-β bands (10– 15 Hz) during motor preparation occurred as distinct dynamics of motor networks across the recovery process. The longitudinal profile of these network dynamics paralleled recovery, suggesting the contribution of the interhemispheric pathway to the increased activity of the ipsilesional sensorimotor cortex and functional recovery. However, this mathematical analysis-based hypothesis awaits biological validation.

Recently, pathway-specific manipulation using chemogenetic or optogenetic tools has been developed not only in model animals such as rodents but also in primates to analyse the causal role of target pathways^11,12^. Here, we blocked the interhemispheric pathway from the contralesional to ipsilesional PM to demonstrate its contribution to dexterous hand movements during recovery in macaque monkeys following lesioning. We also recorded the activity of the sensorimotor cortices with ECoG to study the change in network dynamics caused by the manipulation of the interhemispheric pathway.

## Blockade of the unidirectional interhemispheric pathway using DREADDs

We combined double viral vector intersectional technology^11,13–15^ with Designer Receptors Exclusively Activated by Designer Drugs (DREADDs)^16,17^ for the reversible and unidirectional blockade of the interhemispheric pathway. First, we trained two macaque monkeys to perform a reach and grasp task with their right forelimb (Fig. 1a–b). After the initial training was completed, we injected an anterograde vector (AAV1-EF1α-DIO-hM4D(Gi)-mCherry) into the left PM (mainly in its dorsal part, PMd) and a retrograde vector (AAV2retro-CAGGs-Cre) into the right PM (mainly in the PMd) so that commissural neurons from the left to right PM expressed hM4Di and mCherry (Fig. 1c, Extended Data Fig. 1a). Postmortem anti-RFP immunohistochemistry showed the labelling of the cell bodies of pyramidal neurons in the superficial layers of the left PM, their axons in the corpus callosum, and their terminal axons and buttons in the superficial layer of the right PM (Fig. 1d–h). Note that there were no labelled cell bodies in the right PM, suggesting that unidirectional blockade had been achieved. Conversely, a small number of collateral fibres were detected in the bilateral putamen (Extended Data Fig. 1b), although the density of the labelled fibres was quite low compared with that in the right PM. In addition, we chronically implanted a 28-channel subdural ECoG electrode array on each side of the PM, M1, and primary sensory cortex (S1) for longitudinal recording (Extended Data Fig. 1a).

**Fig. 1.**
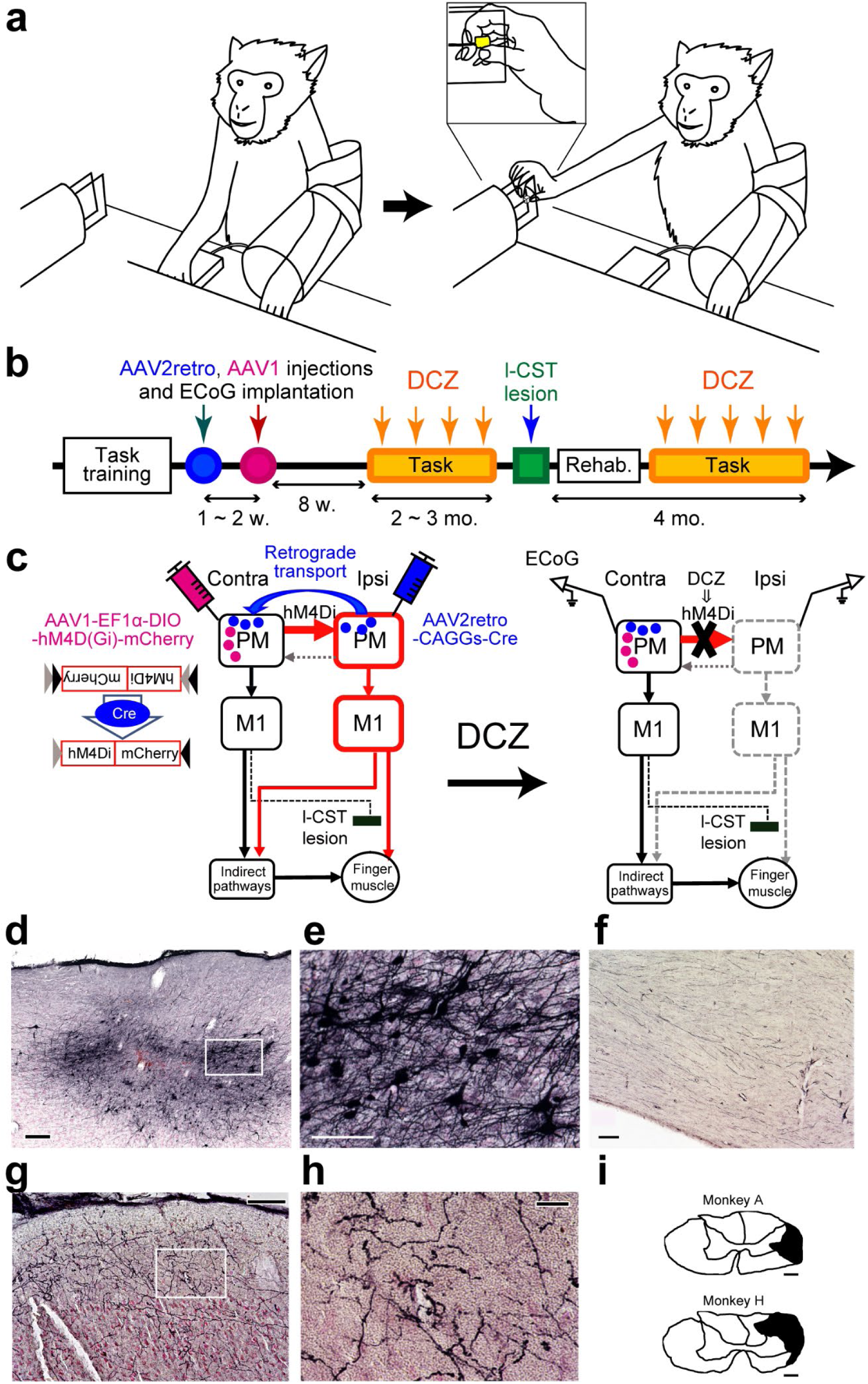
Unidirectional blockade of the interhemispheric pathway using DREADDs. **a**, Reach and grasp task. **b**, Experimental schedule. **c**, Schematic diagrams of vector injections and the mechanism of unidirectional blockade of the interhemispheric pathway. **d**, Representative RFP-labelled cells in the contralesional (left) PM of Monkey A. Rectangle indicates the area shown in ***e***. Scale bar, 200 µm. **e**, High magnification view of ***d***. Scale bar, 100 µm. **f**, RFP-labelled fibres in the corpus callosum of Monkey A. Scale bar, 100 µm. **g**, RFP-labelled fibres in the ipsilesional (right) PM of Monkey A. Rectangle indicates the area shown in ***h***. Scale bar, 100 µm. **h**, High magnification view of ***g***. Scale bar, 20 µm. **i**, Extent of the l-CST lesions. Lesions are indicated by black areas. Scale bar, 1 mm. Rehab., rehabilitation.

## Unidirectional blockade of the interhemispheric pathway affects recovered hand movements

We made a surgical lesion involving the right dorsolateral funiculus at the middle cervical cord (C4– C5) (Fig. 1i). We evaluated performance in the reach and grasp task according to the success rate of precision grip and movement kinematics before lesioning and during recovery. In the prelesional state, the monkeys could retrieve the food pieces with a precise grip, using just the index finger and the pad of the thumb, with a 100% success rate. We injected a DREADD agonist, deschloroclozapine (DCZ)^18^, systemically to block the interhemispheric pathway; however, no significant effect was observed on the success rate of precise grip movements or movement velocity (Fig. 2a, Extended Data Fig. 2a).

**Fig. 2.**
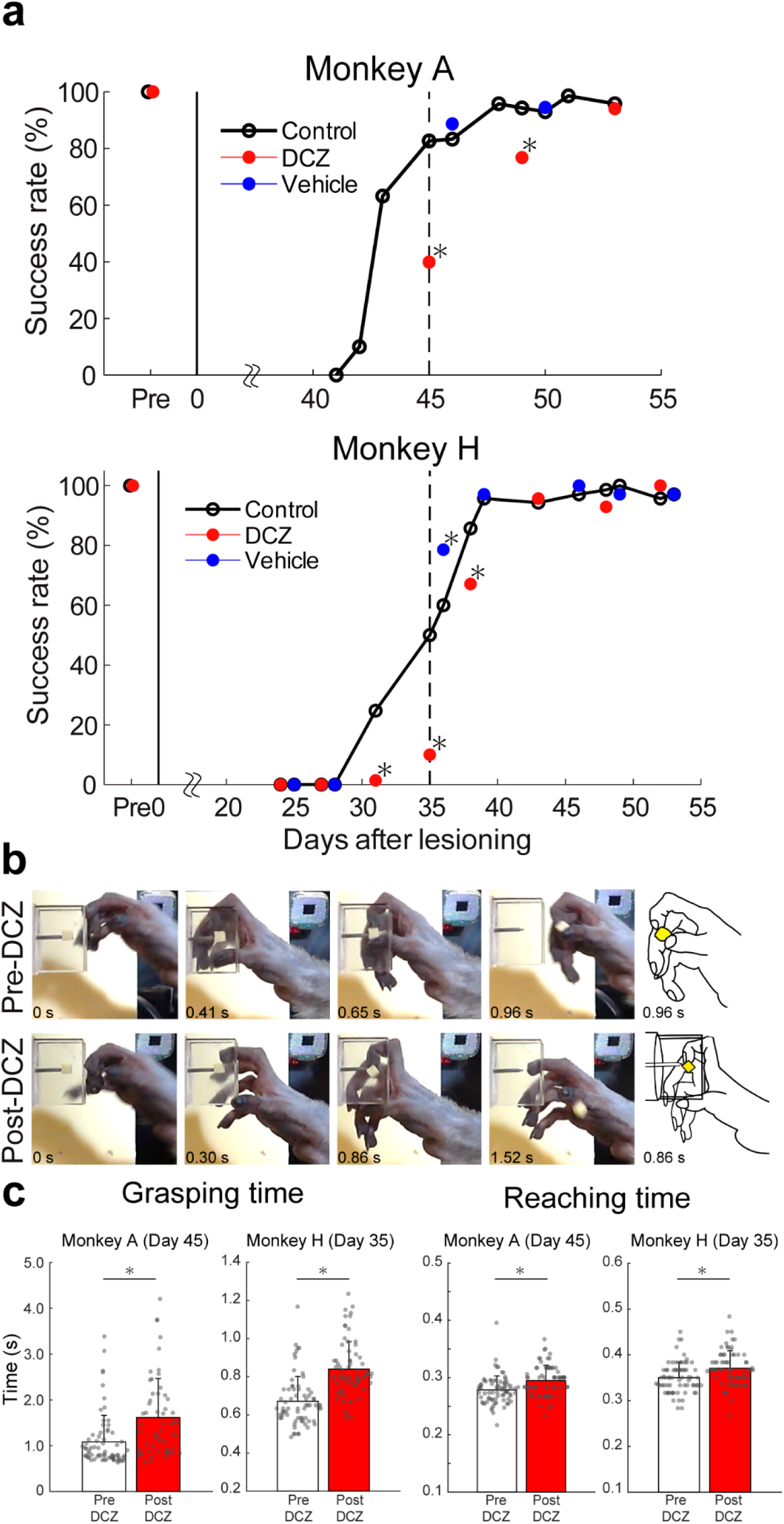
Effects of unidirectional blockade of the interhemispheric pathway on recovered hand movements. **a**, Success rate of precision grip before (open circles) and after (red circles) DCZ administration and after vehicle administration (blue circles). The dotted lines indicate the days when the results of the reaching and grasping times shown in ***c*** were obtained. \**P* < 0.05 (Pearson’s *χ*^2^ test). **b**, Examples of food retrieval behaviour before (upper figures) and after (lower figures) DCZ injection; 45 days after lesioning in Monkey A. **c**, Grasping and reaching time before (white bars) and after (red bars) DCZ administration; 45 days after lesioning in Monkey A (Pre-DCZ: 70 trials, Post-DCZ: 56 trials), and 35 days after lesioning in Monkey H (Pre-DCZ: 70 trials, Post-DCZ: 69 trials). Error bars indicate standard deviation. \**P* < 0.05 (Wilcoxon rank-sum test; Grasping time: *P* = 5.1 × 10^-6^ [Monkey A], 1.3 × 10^-11^ [Monkey H]; reaching time: *P* = 8.3 × 10^-5^ [Monkey A], 7.3 × 10^-4^ [Monkey H]).

Just after lesioning, the monkeys could not move their hands at all; however, they gradually recovered with rehabilitative training. At 1–2 months after lesioning, the monkeys were able to retrieve the food pieces with their fingertips, and the success rate was improved. When we administered DCZ during this period, the recovered dexterous hand movements were impaired and the success rate dropped in both monkeys (Fig. 2a–b); grasping time was also significantly prolonged (Fig. 2c). Machine-learning based analysis of finger trajectories revealed that the monkeys had difficulty in inserting their thumb into the slit (Fig. 2b, Extended Data Fig. 2b–d). In addition, Monkey A failed to retrieve the food pieces and tried to grasp them repeatedly, probably due to weakness of the opposing muscles (Supplementary Video 1). Conversely, the trajectory of the index finger was not different between before and after DCZ administration (Extended Data Fig. 2b).

These results show that the interhemispheric pathway from the contralesional to ipsilesional PM contributes to motor function during recovery, especially the dexterity of the thumb such as opposability. We evaluated motor function until 4 months after lesioning; however, impairment by DCZ blockade was not observed after recovery was accomplished.

## Interhemispheric connectivity is blocked unidirectionally by DREADDs

We previously showed that connectivity calculated with GC^10^ from the contralesional to ipsilesional PM (mainly the PMd) is increased during recovery after l-CST lesioning. In this study, we also recorded ECoG longitudinally and calculated GC between the electrodes during the reach and grasp task. We compared connectivity between before and after DCZ administration to confirm that our manipulation could block interhemispheric connectivity. We divided the recovery stage into the ‘early’ and ‘late’ periods, and the border between ‘early’ and ‘late’ was termed the ‘full recovery day’, which was defined as the experimental day on which the success rate of precision grip first reached 100% after lesioning. During the early recovery stage, GC from the contralesional to ipsilesional PMd in the low frequency band (10–15 Hz) around movement onset (−0.1 – 0.1 s) was decreased after DCZ administration (Fig. 3a–b, Extended Data Fig. 3). We combined the data from both monkeys and found a significant difference in the GC value before and after using DCZ (Fig. 3d). Conversely, GC in the opposite direction (from the ipsilesional to contralesional PMd) was not significantly changed by DCZ administration (Extended Data Fig. 3c–d). These results suggest that our method successfully blocks the connectivity of the target pathways in a unidirectional manner.

**Fig. 3.**
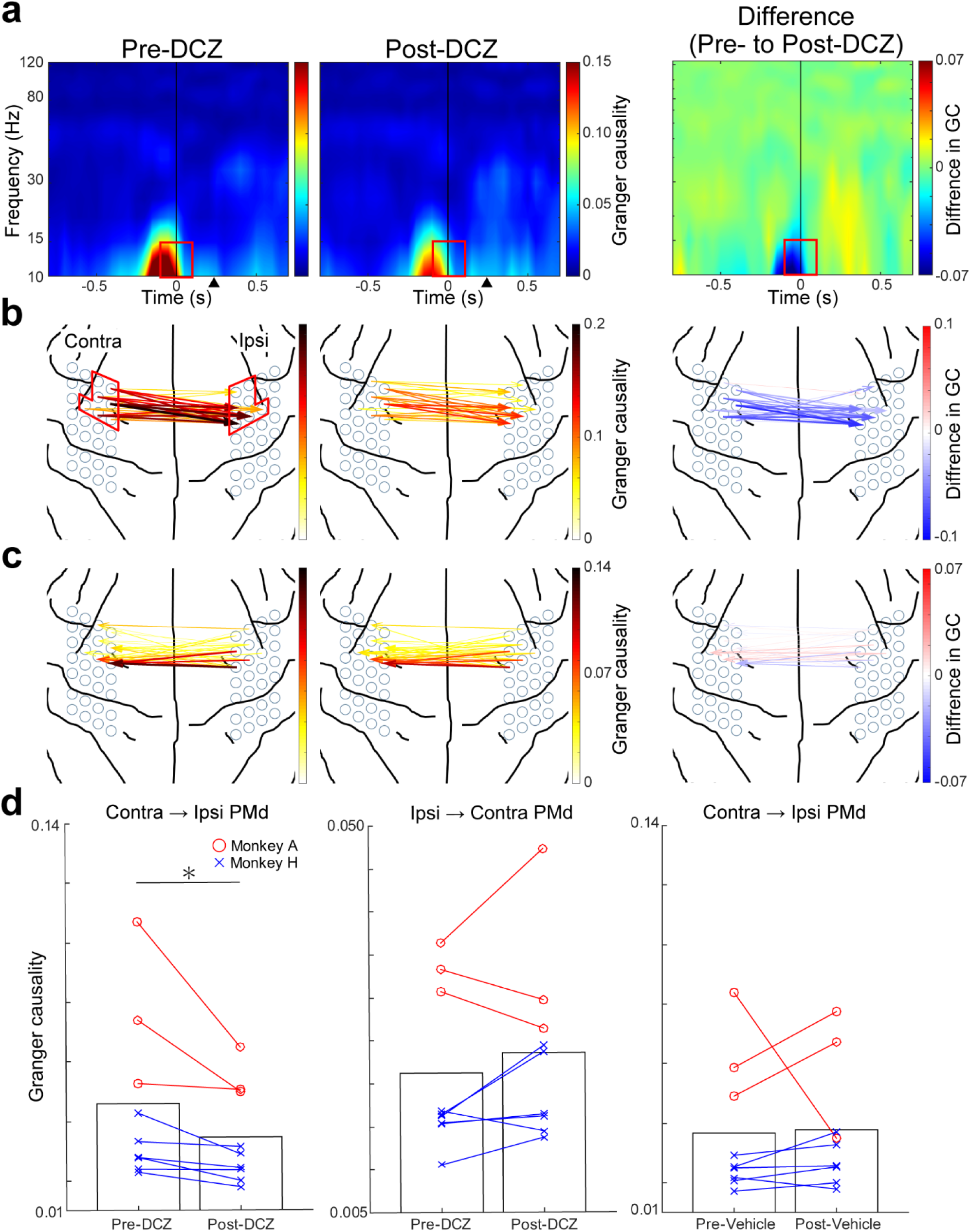
Interhemispheric connectivity is unidirectionally blocked by DREADDs. **a**, Averaged time-frequencygram of GC from the contralesional to ipsilesional PM, particularly PMd (7 × 7 channels; the area surrounded by red lines in the left panel of ***b*** before (left figure) and after (middle figure) DCZ administration in the early recovery stage of Monkey A. Right figure shows the difference in GC between before and after DCZ administration. Time 0 (black line) indicates movement onset. Red rectangles indicate the time frequency ROI that was used for the calculations shown in ***b–d***. Black triangles at the bottom indicate the onset of grasping, which we defined as the timing when the monkey’s finger first touched the slit. **b**, Anatomical dimension of GC from the contralesional to ipsilesional PMd (7 × 7 channels, encircled in red lines) in the low-frequency band around movement onset (ROI shown in ***a***), before (left) and after (middle) DCZ administration and their difference (right). Each arrow indicates the directionality of GC. Its colour and width indicate the strength of GC. **c**, Anatomical dimension of GC from the ipsilesional to contralesional PMd before (left) and after (middle) DCZ administration and their difference (right). **d**, Averaged value of GC in the ROI (shown in ***a***) in the early recovery stage of both monkeys (A and H). Each line plot indicates the result of each experimental day in the early recovery stage (*n* = 9 days). Bars indicate the averaged value of all experimental days in both monkeys. Each figure shows the GC from the contralesional to ipsilesional PMd with DCZ (left) or vehicle (right) and GC in the opposite direction with DCZ (middle), respectively. \**P* < 0.05 (Wilcoxon sign-rank test; *P* = 0.0039 [left], 0.25 [middle], 0.35 [right]).

## Interhemispheric facilitation during recovery

We investigated the longitudinal change in the activity of the ipsilesional PM during blockade of the interhemispheric pathway. We performed time-frequency analysis of ECoG activity during the task and compared the activity between before and after DCZ administration. In the intact state, the activity of the ipsilesional PM in the low frequency band (7–9 Hz) around movement onset (−0.2 – 0.1 s) was increased by DCZ administration (Fig. 4a–b). In the early recovery stage, it was decreased by DCZ (Fig. 4c–d). Furthermore, in the late recovery stage, DCZ administration increased activity again, as in the intact state (Fig. 4e–f). Vehicle had no effect on activity in any stage (Extended Data Fig. 4). These results indicate that the interhemispheric pathway inhibits the activity of the opposite PM in the intact state, while this pathway facilitates the ipsilesional PM and contributes to recovery in the early recovery stage (Fig. 4g).

**Fig. 4.**
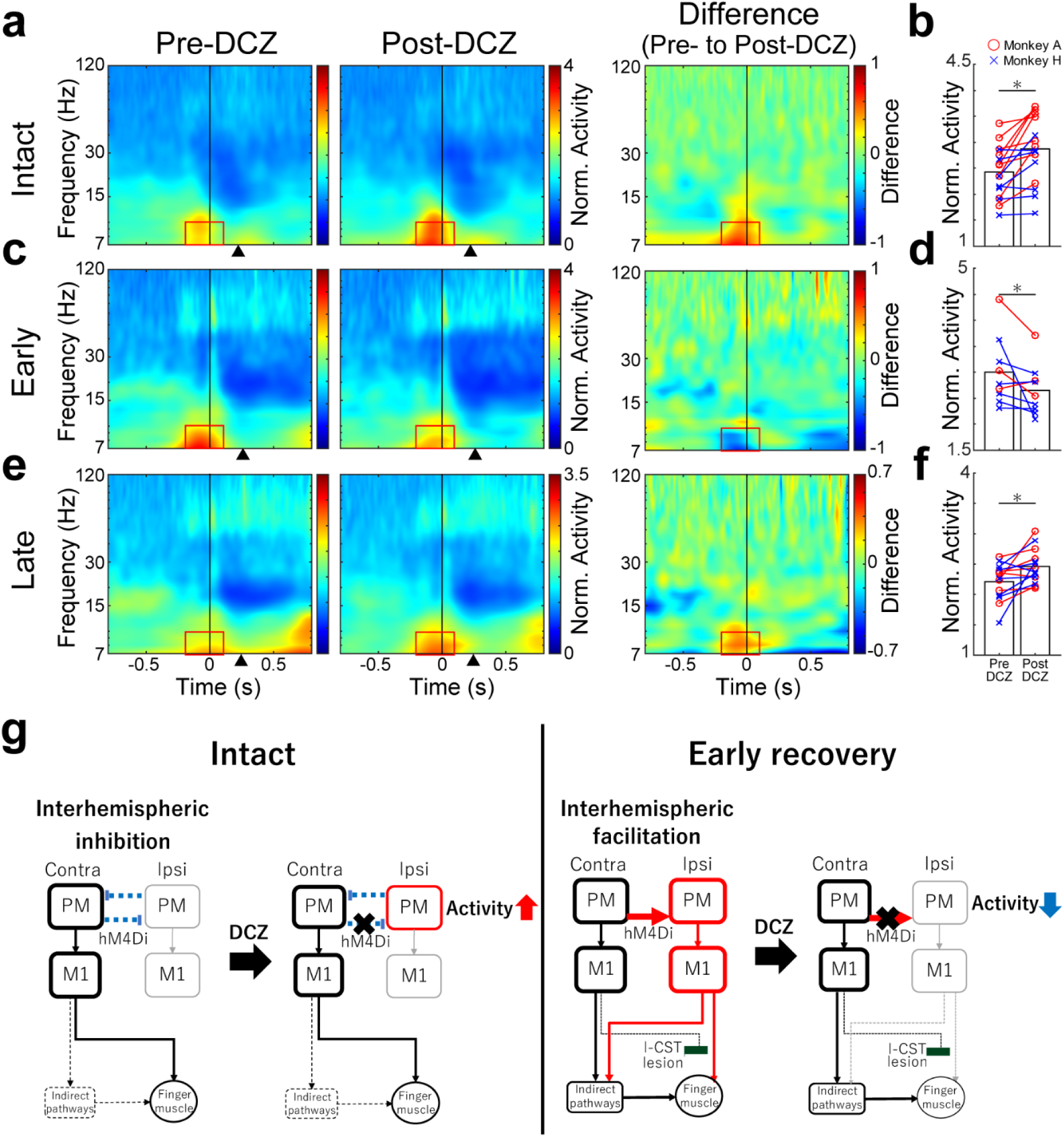
Interhemispheric facilitation during recovery. **a**,**c**,**e**, The average normalized time-frequencygram of brain activity (ECoG) at the ipsilesional PMd (Ch 10) before (left figures) and after (middle figures) DCZ administration in the intact state (**a**, *n* = 16 experiment days including 1,074 trials [Pre-DCZ] and 1,093 trials [Post-DCZ]), early recovery stage (**c**, *n* = 9 days, 591 trials [Pre-DCZ] and 525 trials [Post-DCZ]), and the late recovery stage (**e**, *n* = 14 days, 970 trials [Pre-DCZ] and 972 trials [Post-DCZ]) of both monkeys. Right figures show the difference in activity between before and after DCZ administration. Time 0 (black line) indicates movement onset. Red rectangles indicate the time frequency ROI that was used for the calculations shown in ***b****,**d**,**f***. Black triangles indicate the onset of grasping. **b**,**d**,**f**, Averaged value of normalized brain activity (ECoG) at the ipsilesional PMd in the ROI in the intact state (**b**), early recovery stage (**d**), and late recovery stage (**f**) of both monkeys (A and H). Each line plot indicates the result of each experimental day and bars indicate the averaged value of all experimental days in both monkeys. \**P* < 0.05 (Wilcoxon sign-rank test, *P* = 9.4 × 10^-4^ [***b***], 0.039 [***d***], 0.0067 [***f***]). **g**, Diagram showing the hypothesis drawn from the present findings. Red lines indicate the activated areas and pathways. Dotted lines indicate the suppressed areas and pathways.

In addition, we performed electrical stimulation with the subdural electrodes on the bilateral sensorimotor cortices and observed muscle twitches in the ipsilesional forelimb. In the intact state, the muscle twitches of the ipsilesional forelimb were barely observed when the ipsilesional electrodes were activated. However, during the early recovery stage, electrical stimulation on the ipsilesional side induced twitches in the ipsilesional forelimb muscles (Extended Data Fig. 5a–b), suggesting the involvement of the ipsilesional sensorimotor cortex in the control of affected hand movements.

We also recorded cortico-cortical evoked potentials (CCEPs)^19^ at the ipsilesional PM by stimulating the contralesional PM. The amplitude of CCEPs was increased during the early recovery stage (Extended Data Fig. 6).

These results supported our conclusion that the interhemispheric pathway from the contralesional PM facilitates the activity of the ipsilesional sensorimotor cortex, and the pathway from the ipsilesional motor cortex to the ipsilesional hand muscles is upregulated and supports recovery during the early stage after lesioning (Fig. 4g).

## Discussion

The early recovery stage after lesioning is critical because a previous study showed that the lack of training during the first month after l-CST lesioning has a severe impact on recovery^20^. In the present study, we demonstrated that signal flow from the contralesional to ipsilesional PM caused interhemispheric facilitation of the ipsilesional sensorimotor cortex and contributed to recovery during the early stage after l-CST lesioning by using a state-of-the-art intersectional pathway-specific blocking technique.

In the intact state, blockade of the interhemispheric pathway did not affect motor performance in the reach and grasp task, showing that neither the interhemispheric pathway nor ipsilateral motor cortex contributed to hand movements. Time-frequency analysis suggested that the interhemispheric pathway from the left PM inhibited the activity of the right PM in the intact state, which is consistent with previous reports^21,22^. Interestingly, the present results suggest that the effect of the interhemispheric pathway changes from inhibition to facilitation and the ipsilesional sensorimotor cortex becomes involved in the control of dexterous hand movements in the early recovery stage after lesioning. Commissural fibres originate from glutamatergic excitatory pyramidal neurons and exert excitatory and inhibitory effects on the pyramidal neurons in the opposite sensorimotor cortex^23,24^. Therefore, the disinhibition observed in the early recovery stage could be due to changes in the activity of inhibitory interneurons at the ipsilesional PM or the postsynaptic effect of GABA from inhibitory to excitatory due to changes in Cl^-^ transporters, as observed during the early developmental stage or in some pathological conditions^25^, but the detailed mechanisms still remain unclear.

More interestingly, DCZ blockade had no effect on behaviour in the late recovery stage, and time-frequency analysis showed the effect of DCZ in this period was the same as in the intact state. These results suggest that the action of the interhemispheric pathway returned to inhibition after full recovery. These findings were consistent with previous studies also suggesting that the contribution of the interhemispheric pathway and ipsilesional motor cortex was more obvious in the early stage than in the chronic stage^9,10^. Our recent neuroanatomical study demonstrated that late stage recovery is supported by reorganization of the CST including the sprouting of axons originating from the contralesional and ipsilesional motor cortices in the grey matter both caudal and rostral to the lesion, including a re-direction of the CST to hand motoneurons^26^. In addition, other descending motor pathways such as propriospinal and reticulospinal tracts remained intact in our lesion model, and these pathways might also contribute to the recovery of hand movements.

In the clinical setting, many studies have focused on the role of the interhemispheric pathway in patients with a CST lesion such as stroke^27–29^. However, it is still controversial whether the interhemispheric pathway and motor cortex on the intact side support or disrupt recovery^30,31^. It is thought that the results depend on lesion size, recovery stage, comorbidities, etc., but these are difficult to control in clinical studies. One of the advantages of our research was that we could follow the whole process longitudinally before and after a particular lesion. In addition, pathway-specific manipulation using chemogenetic techniques is one of the best approaches to reveal the causal relationship between a target pathway and its functions. Furthermore, since it enabled us to block the pathway reversibly and repeatedly, we could investigate its role longitudinally in different phases with a minimum number of animals.

There are some technical concerns about our pathway-specific blockade method. First, although DCZ is a high-affinity agonist for DREADDs, it could still have off-target effects on endogenous receptors. However, it is unlikely that our behavioural results were due to off-target effects because we examined another monkey without DREADD expression and found that DCZ did not affect its behaviour in the early recovery stage (Extended Data Fig. 7a). Second, although vector expression was quite selective to the interhemispheric pathway, some collateral fibres do project to the bilateral putamen because intratelencephalic neurons have collateral projections to the striatum in rodents^32,33^. However, it is unlikely that blocking such collateral fibres led to the current results because the density of the labelled fibres was quite low compared with that in the ipsilesional PM. In addition, we performed a partial callosotomy between the PMs during recovery after l-CST lesioning and obtained similar behavioural results (Extended Data Fig. 7), suggesting that commissural fibres were the main contributors to our results.

Our findings demonstrated that the contribution of the interhemispheric pathway was observed in the early recovery stage. In the clinical setting, the enhancement of functional recovery in the early stage with neurorehabilitation is important for patients to increase their activities of daily living and motivation. Otherwise, poor functional recovery in the early stage results in disuse and muscle atrophy in the late stage. Our results offer new insights on a target pathway for neuromodulation therapy to enhance recovery during the early phase after lesioning of the CST.

## Methods

### Subjects

We used two Japanese macaque monkeys (*Macaca fuscata*; Monkey A [5 years old, 7 kg, male] and Monkey H [7 years old, 8 kg, female]) in the pathway-specific blockade experiments. We used another two monkeys (Monkey I [6 years old, 7 kg, female] and Monkey K [6 years old, 6 kg, female]) for the callosotomy experiments (Extended Data Fig. 7). All experimental procedures and animal care were performed in accordance with the ILAR’s Guide for the Care and Use of Laboratory Animals and were approved by the Committee for Animal Experiments at the Graduate School of Medicine in Kyoto University, Japan.

### Experiments with pathway-specific blockade

#### Surgery

We performed all surgical procedures described below under general anaesthesia. First, we anesthetized the animals with ketamine hydrochloride (10 mg/kg, i.m.) and xylazine (1 mg/kg, i.m.) and then intubated them and maintained anaesthesia with isoflurane (1–2%) inhalation. We monitored heart rate, blood pressure, peripheral capillary oxygen saturation, body temperature, and end-expiratory carbon dioxide pressure during surgery. We administered dexamethasone (0.825 mg/kg, i.m.), analgesics (butorphanol [0.2 mg/kg, i.m.], ketoprofen [1 mg/kg, i.m.], or diclofenac [12.5 mg, anally]), and antibiotics (ampicillin sodium [200 mg, i.m.], ceftriaxone [50 mg, i.m.], or cefazolin [100 mg, i.m.]) during and after surgery.

#### Implantation of steel tubes for head fixation

First, we performed magnetic resonance imaging of the brain under anaesthesia with ketamine hydrochloride (10 mg/kg, i.m.) and xylazine (1 mg/kg, i.m.) and then intubated the animals and maintained anaesthesia with isoflurane (1–2%) inhalation using a 3T magnetic resonance imaging scanner (Verio; Siemens, Washington, DC, USA) to decide the location of plastic screws, steel tubes, and craniotomy.

We mounted two steel tubes on the monkey’s head for fixation during the behavioural task. First, we made a skin incision on the head. We peeled off the soft tissue and a part of the temporal muscles to expose the skull. We attached small plastic screws to the skull as anchors. We mounted two steel tubes in parallel over the frontal and occipital lobes. Finally, we covered the skull, screws, and steel tubes completely with acrylic resin.

#### Injection of viral vectors and implantation of ECoG electrodes

To block signal transmission through the interhemispheric pathway from the contralesional (left) to ipsilesional (right) PM, we injected AAV1-EF1α-DIO-hM4Di-mCherry (1.5 × 10^12^ vg/mL [Monkey A] and 2.2 × 10^12^ vg/mL [Monkey H]) into the contralesional PM and AAV2retro-CAGGS-Cre (6.0 × 10^12^ vg/mL [Monkey A] and 5.3 × 10^12^ vg/mL [Monkey H]) into the ipsilesional PM. We prepared all vectors as described previously^34^. A transferred plasmid, pAAV-hSyn-hM4D(Gi)-mCherry was a gift from Bryan Roth (Addgene plasmid # 50475; http://n2t.net/addgene:50475; RRID:Addgene_50475). We exposed the PM, M1, and S1 by removing the acrylic resin, craniotomy, and incision of the dura mater. We injected six tracks in the PMd (the area between the spur of the arcuate sulcus and superior precentral sulcus) and two to three tracks in the ventral part of the PM (lateral to the spur of the arcuate sulcus). The distance between the tracks was 2.0–2.5 mm (Extended Data Fig. 1). To spread the vector solution to a large area of the cortex, we adopted a convection-enhanced delivery technique^35,36^, which enabled the injection of a large volume of solution at a controlled high rate. Specifically, we injected 6.05 μL vector at each track in two steps. We injected the first 1.25 μL at 0.25 μL/min, and injected the remaining 4.80 μL at 0.60 μL/min by using a 25-μL Hamilton syringe (1702SN, flat tip; Hamilton Company, Reno, NV, USA) with a 32-gauge injection needle. We fitted a fused silica capillary (450-μm outside diameter) to create a 1 mm ‘step’ away from the needle tip to reduce reflux. We inserted the needle 1.5 mm from the surface of the cortex. Before and after injection, we waited for 5 and 10 min, respectively.

After the injection, we implanted a platinum ECoG array comprised of 28 channels (7 × 4 grid) electrodes on a parallelogram-shaped silicon sheet (23 × 11.5 mm) covering the PM where the injections were made, M1 hand area, and S1. The diameter of each electrode was 2 mm and the inter-electrode distance was 3 mm (the distance between the electrodes on the M1 and S1 was 4 mm). There were four other electrodes on the opposite surface of the sheet for reference. We performed the surgeries one side at a time in two separate days.

#### Lesioning of the l-CST

We lesioned the l-CST at the right middle cervical cord (C4–C5) as described previously^10,37^. Briefly, we exposed the dorsal surface of the spinal cord by laminectomy of the C3 and C4 vertebrae and incision of the dura mater. We transected the dorsal part of the lateral funiculus at the border between the C4 and C5 segments from the dorsal root entry zone ventrally to the level of the lateral ligaments by using a pair of fine forceps. We extended the lesion ventrally at the most lateral part of the lateral funiculus. We closed the opening of the dura mater by using artificial dura mater (Gore-Tex membrane; WL Gore & Associates, Flagstaff, AZ, USA), and we sutured the muscles and skin with absorbable sutures and silk, respectively.

#### Behavioural testing

First, we trained the monkeys to sit in a monkey chair with their heads fixed in a stereotaxic frame attached to the chair and with their left arm fixed gently in the arm holder (Fig. 1a). The monkeys were required to push a button on a board attached to the chair with their right hand for more than 2 s. Then, a cube of sweet potato (6 × 6 × 6 mm^3^) was presented between a vertical slit (8 mm width), and the monkeys could retrieve it with their thumb and index finger. Signals from the button allowed us to determine the timing of movement onset. We used two digital video cameras (120 frames/s) to record and analyse the reach and grasp task; one from the left and another from the upper side of the monkeys. Since the monkeys could not perform the reach and grasp task just after l-CST lesioning, we performed intensive rehabilitation for 5–6 days per week with manual assistance of movements and with larger pieces of sweet potatoes and apples to promote rehabilitation until they were able to perform the task.

#### ECoG recording during behavioural testing

While the monkeys performed the reach and grasp task, we recorded brain activity on the surface of the bilateral sensorimotor cortices using the ECoG electrodes. We recorded the signals at a sampling rate of 2,000 Hz. We extracted ECoG signals using multichannel amplifiers with 0.3-Hz high-pass and 7,500-Hz low-pass analogue filters.

#### Administration of DCZ

We used DCZ (HY-42110; MedChemExpress, Monmouth Junction, NJ, USA), a high-affinity and selective agonist for DREADDs^18^, to block the interhemispheric pathway. We dissolved DCZ in 2% dimethyl sulfoxide in saline (vehicle) to a final concentration of 1 mg/mL. Before DCZ administration, the monkeys performed 70 trials of the reach and grasp task as the Pre-DCZ session. After the Pre-DCZ session, we administered DCZ (100 µg/kg) intramuscularly and the monkeys performed 70 trials of the task as the Post-DCZ session. The Post-DCZ session was started at 30 min after DCZ administration and was accomplished by 60 min after administration. We administered DCZ or vehicle once or twice per week during the prelesional period, for 2–3 months before lesioning, and for 4 months in the postlesional period.

#### Behavioural data analysis

We calculated the success rate of precision grip during the reach and grasp task as described previously^11,13^. We identified three types of error: ‘drop error’, ‘wandering error’, and ‘precision grip error’. For drop errors, the monkeys dropped the food and failed to eat it. We defined wandering errors as trials in which both fingers touched the food, but the monkeys released it and tried to pick it up again. If the monkeys could not pinch the food with their index finger and the pad of the thumb, we defined these trials as precision grip error trials, although they were able to eat the food. In this study, we evaluated success rates by removing these errors. We compared the success rates of the Pre- and Post-DCZ sessions on each day by using Pearson’s χ^2^ test (two-sided).

We calculated the reaching and grasping times in each trial. We defined reaching time as the time from movement onset to when the monkey’s finger first touched the slit, while grasping time was the time during which the monkey’s fingers were inside the slit. We removed drop error trials from the analysis. We tested the significance of the difference in the reaching and grasping times between the Pre- and Post-DCZ sessions by using the Wilcoxon rank-sum test (two-sided).

We tracked the position of the thumb and index finger recorded from the left side by using machine-learning software (DeepLabCut)^38^. Briefly, we selected 500 frames randomly from two videos that recorded the Pre- and Post-DCZ sessions on the same experimental day that we used to label the tips of the thumb and index finger, food, and upper and lower right edges of the slit. We trained the deep learning network over 500,000 iterations and fed it both videos to track the tips of the fingers during the task. We extracted the coordinates of all labels using MATLAB (MathWorks, Natick, MA, USA). We removed the drop error trials and plotted the trajectories of the thumb and index finger in each trial (Extended Data Fig. 2b) at the timing just before the monkeys retrieved the food. We plotted the positions of the thumb and index finger (Extended Data Fig. 2c) at the timing of retrieval, which we defined as the timing at which the *x* coordinate of the food first exceeded the baseline position by a threshold of 5 standard deviations. We calculated the distance of the thumb tip from the slit entrance at the timing when the monkeys retrieved the food as an index for the dexterity of the thumb, which indicates how well they could insert the thumb into the slit. We tested the significance of the difference in the distance between the Pre- and Post-DCZ sessions by using the Wilcoxon rank-sum test (two-sided) (Extended Data Fig. 2d).

#### Dynamics of cortical activity

To evaluate the dynamics of cortical activity, we performed time-frequency analysis of the ECoG data as described previously^10^. Briefly, we removed the 60-Hz line noise from the raw ECoG data using the MATLAB FieldTrip toolbox^39^. We downsampled the data four times, resulting in a sampling rate of 500 Hz. We then aligned the ECoG signals from each channel using the timing of reaching onset, which we defined as the timing when the monkeys left the button (Time 0; see Fig. 4a). The drop error and wandering error trials were removed from the analysis. By using the MATLAB FieldTrip toolbox, we quantified the dynamics of cortical activation in each channel by the time-frequency representation generated by the Morlet wavelet transform method at 116 different centre frequencies (5–120 Hz) with the half-length of the Morlet analysing wavelet set at the coarsest scale of five samples. We normalized each time-frequency representation value by the baseline value (mean time-frequency representation value at the corresponding frequency during the resting period from - 1.5 – −1 s).

We set the time frequency region of interest (ROI) in the low frequency band (7–9 Hz) around movement onset (−0.2 – 0.1 s), where the activity was the most prominent (Fig, 4a,c,e), and calculated the mean normalized time-frequency representation of the ROI in each session of each experimental day. We divided the experimental period into three groups: ‘intact’ (pre-lesion), ‘early’ recovery stage, and ‘late’ recovery stage. We termed the border between early and late as the ‘full recovery day’, which we defined as the experimental day on which the success rate of precision grip first reached 100% (day 57, Monkey A; day 49, Monkey H). We tested the significance of the difference in the mean activity of the ROI between the Pre- and Post-DCZ sessions in each recovery stage (intact, *n* = 9 for Monkey A, *n* = 7 for Monkey H; early, *n* = 3 for Monkey A, *n* = 6 for Monkey H; late, *n* = 7 for Monkey A, *n* = 7 for Monkey H) by using the Wilcoxon sign-rank test (two-sided).

#### Dynamics of corticocortical connectivity

We quantified the dynamics of corticocortical connectivity by spectral GC^40^, as described previously^10^. Briefly, we downsampled the data eight times, resulting in a sampling rate of 250 Hz. We then aligned the ECoG signals from each channel at reaching onset. We removed the drop error and wandering error trials from the analysis. Between the signals from two channels (56 × 56 channels), we calculated in-trial spectral GCs from each session, where each GC represents a unidirectional connectivity from one channel to another, during times between −1.5 and 1 s (125 time points), and across frequencies between 5 and 120 Hz.

We used three preparation steps for spectral GC calculation:

1. Preprocessing: we performed detrending, temporal normalization, and ensemble normalization to achieve local stationarity of the data^41^;
2. Window length selection: we set the length and step size of the sliding window for segmentation as 150 ms and 20 ms, respectively;
3. Model order selection: we set the model order, which is related to the length of the signal in the past that is relevant to the current observation, as 10 samples (equivalent to 10 × 4 = 40 ms of history) in both monkeys, following a previous study^10^.

After GC calculation, we averaged all combinations of GC from the contralesional PMd to ipsilesional PMd (7 × 7 channels, Fig. 3b) or the opposite direction. We set the time frequency ROI in the low frequency band (10–15 Hz) around movement onset (−0.1 – 0.1 s) where GC was the most prominent in both monkeys (Fig. 3a, Extended Data Fig. 3a), and calculated the mean GC of the ROI in each session of each experimental day. We tested the significance of the difference in the mean GC of the ROI between the Pre- and Post-DCZ sessions in each experimental day during the early recovery stage (*n* = 3 for Monkey A, *n* = 6 for Monkey H) by using the Wilcoxon sign-rank test (two-sided).

#### Electrical stimulation

We performed electrical stimulation experiments by using the ECoG electrodes several times before l-CST lesioning and once a week after lesioning. We trained the monkeys to sit on the monkey chair during stimulation under awake conditions. We delivered current pulses (monophasic, 3 pulses with 20 Hz, 3–4 mA, pulse duration: 0.5 ms) 20–30 times with a 2-s interval through a stimulator (Nihon Kohden, Tokyo, Japan). In response to stimulation, we recorded the muscle twitches in the right (affected side) forearm and CCEPs on the hemisphere opposite to the stimulated side. We evaluated the magnitude of muscle twitches in each body part by using the following criteria: 1) no response; 2) invisible muscle twitches; 3) visible muscle twitches without joint movements; and 4) muscle twitches with joint movements (Extended Data Fig. 5). We calculated the baseline to first negative peak amplitude of the CCEPs and tested the significance of the difference in the amplitude between each recovery period by using one-way analysis of variance and *post-hoc* pairwise comparisons with Bonferroni’s correction (Extended Data Fig. 6b).

We also evaluated the change in the amplitude of CCEPs recorded at the ipsilesional PM between before and after intramuscular DCZ administration (100 µg/kg). However, DCZ did not show any significant effect on CCEPs in any recovery stage (data not shown). Possible reasons we considered are as follows.

1. DREADDs including hM4Di are expressed at the surface of cell bodies and axon terminals^12^. As electrical stimulation directly activates Na^+^ channels at the axon hillock and evokes action potentials, hM4Di expressed at the surface of the cell bodies does not affect CCEPs.
2. hM4Di expression at the axon terminals might not be sufficiently high to change CCEPs, although it did affect spontaneous or physiological activity including the event-related activity shown in Fig. 3. One factor was that its expression with the double viral vector technique could be relatively low compared with single vector injection.

#### Histological assessment and anti-RFP immunohistochemistry

After the behavioural experiments were completed, we anaesthetized the monkeys deeply with thiopental sodium (25 mg/kg, i.v.) and perfused them transcardially with 0.1 M phosphate-buffered saline (PBS), followed by 4% paraformaldehyde in PBS.

We extracted and preserved the whole brain and cervical and upper thoracic spinal cord in 4% paraformaldehyde overnight for post-fixation. After saturation with 30% sucrose solution in PBS, we used a freezing microtome (Retoratome; Yamato Kohki Industrial, Saitama, Japan) to make serial sections (40-μm thick) of the brain (coronal) and spinal cord (axial).

We processed the brain sections for anti-RFP immunohistochemistry. Specifically, we incubated the sections with a rabbit anti-RFP antibody (1:2,000; Rockland Immunochemicals, Inc. Boyertown, PA, USA) and then with a biotinylated goat anti-rabbit IgG antibody (1:200; Vector Laboratories, Burlingame, CA, USA). We visualized the immunoreactive signals with diaminobenzidine (1:10,000; Wako, Tokyo, Japan) containing 1% nickel sodium ammonium and 0.0003% H_2_O_2_ in Tris-buffered saline. We counterstained the sections with Neutral Red. We processed the spinal cord sections for Nissl staining with 0.1% cresyl violet to evaluate the size of the l-CST lesion.

### Experiments with callosotomy

We trained Monkey I to perform the reach and grasp task and then lesioned the l-CST at the right C4–C5. Before lesioning and during recovery, we administered DCZ (100 µg/kg) intramuscularly and measured the success rate of precision grip (Extended Data Fig. 7a) to confirm that DCZ by itself did not affect task performance during recovery in a monkey without DREADD expression. We performed a partial callosotomy at the rostral part of the corpus callosum where the callosal fibres connect to the bilateral PMs during the early recovery stage (35 days after l-CST lesioning). Briefly, under the general anaesthesia described above, we performed a craniotomy and incision of the dura mater at the vertex. We accessed the corpus callosum with gentle retraction of the right hemisphere and coagulated the corpus callosum between the bilateral PMs by using bipolar forceps. After callosotomy, we sutured the dura and skin.

We trained Monkey K to perform the task and it underwent the same partial callosotomy without l-CST lesioning to confirm that callosotomy in the intact state did not affect the success rate of the task (Extended Data Fig. 7b).

## Data availability

The data supporting the findings of this study are available from the corresponding authors upon reasonable request.

## Code availability

The codes supporting the findings of this study are available from the corresponding authors upon reasonable request.

## Acknowledgements

We thank Masashi Nakamura and Erika Omae for technical assistance. We thank Bryan Roth for providing the plasmid. This work was supported by a Grant-in-Aid for Scientific Research on Innovative Areas ‘Hyper-adaptability’ to T.I. (Project no. 19H05723), a Grant-in-Aid for Scientific Research from the Ministry of Education, Culture, Sports, Science, and Technology (MEXT) to T.I. (KAKENHI (A) no. 19H01011 and (S) no. 22H04992), Japan Agency for Medical Research and Development (JP18dm0307005 to N.S. and T.I.), and a Grant-in-Aid for Scientific Research from MEXT to H.O. (KAKENHI (A) no. 20H00573) and to R.Y. (KAKENHI (B) no. 21H02798).

## Author’s contribution

M.M., R.Y., and T.I. designed the experiments with contributions from H.O. and R.T. K.K. designed and produced the viral tools. M.M., R.Y., T.K., S.U., Y.S., and H.O. performed the surgeries. M.M., R.Y., T.K., S.U., and Y.S. conducted the behavioural and electrophysiological experiments. M.M., S.U., K.I., H.O., and J.T. conducted the histological and imaging experiments. M.M., R.Y., and S.U. analysed the experimental data. R.T. and T.I. supervised the study. M.M. and T.I. wrote the manuscript with input from all authors.

---

**Extended Data Fig. 1.**
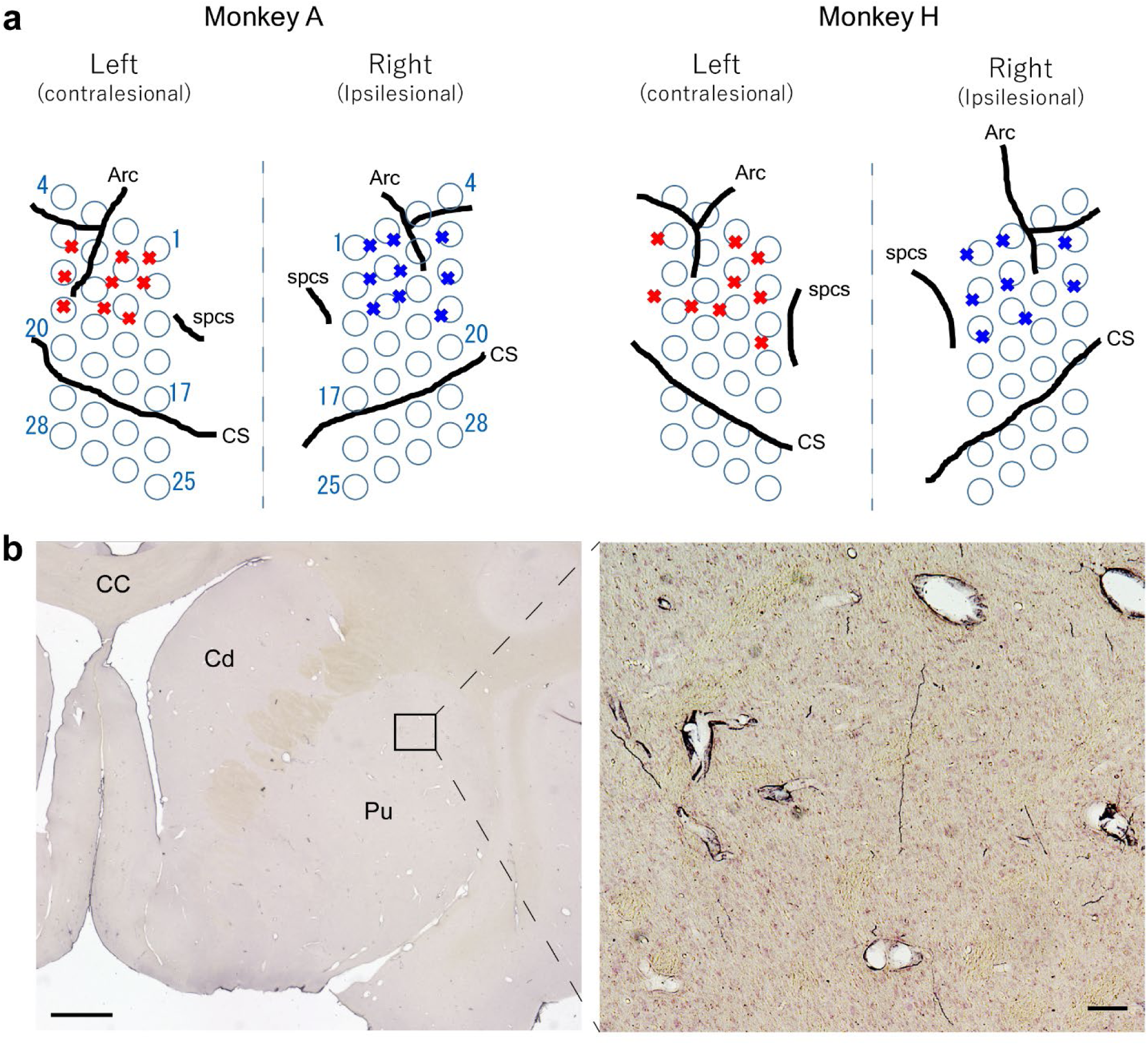
Double viral vector injections for blockade of the unidirectional interhemispheric pathway. **a**, Locations of vector injections and ECoG electrodes in Monkeys A and H. Red and blue crosses indicate the injection sites of AAV1-EF1α-DIO-hM4Di-mCherry and AAV2retro-CAGGS-Cre, respectively. Blue circles indicate the location of the ECoG electrodes. **b**, Representative RFP-labelled fibres in the ipsilesional putamen of Monkey A. Rectangle in the left figure indicates the area shown in the right figure. Scale bar, left: 2 mm, right: 100 µm. Arc, arcuate sulcus; CS, central sulcus; spsc, superior precentral sulcus. CC, corpus callosum; Cd, caudate; Pu, putamen.

**Extended Data Fig. 2.**
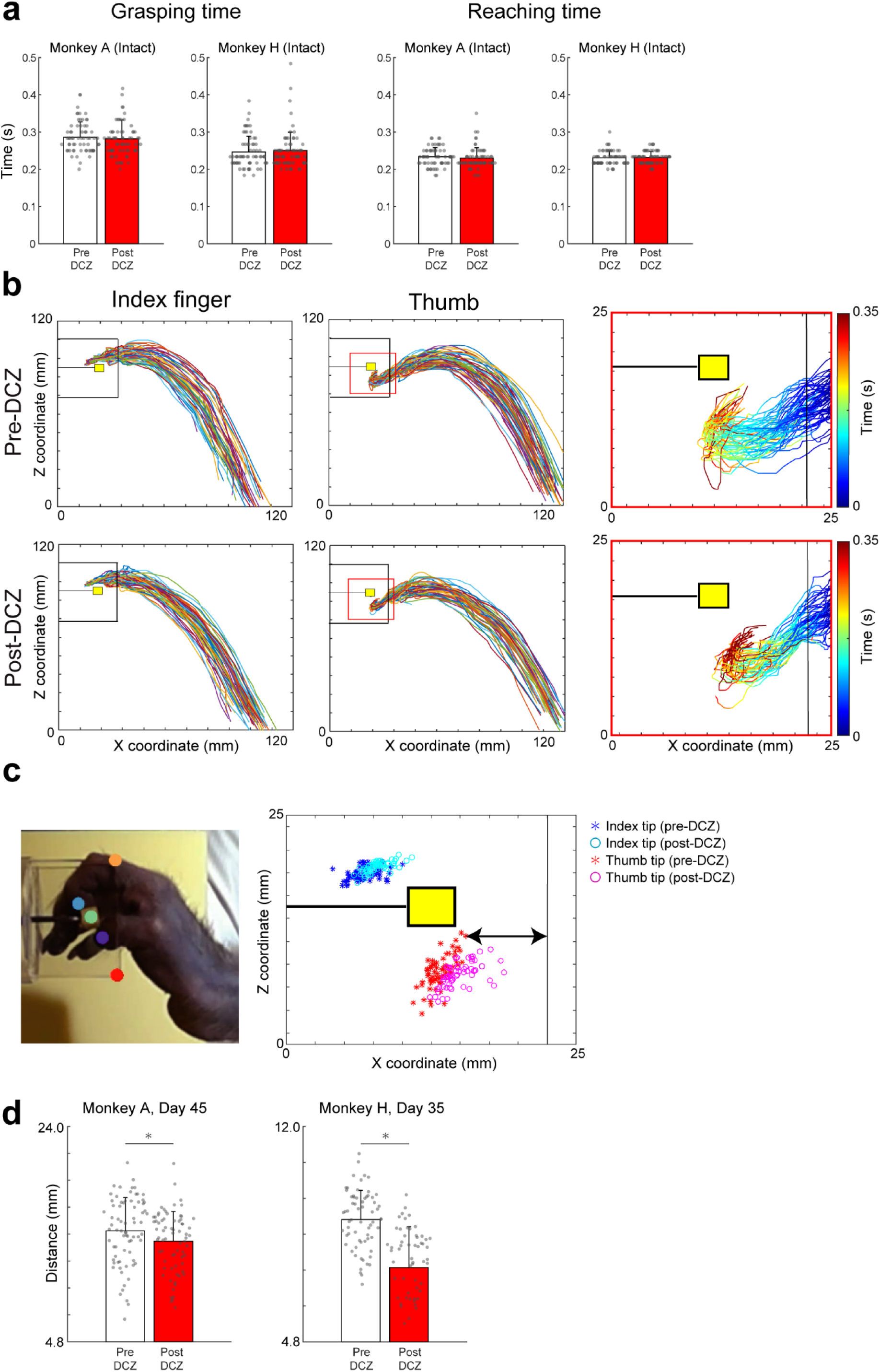
Blockade of the interhemispheric pathway affects thumb dexterity. **a**, Reaching and grasping times before (white bars) and after (red bars) DCZ administration before lesioning in Monkey A (Pre-DCZ: 70 trials, Post-DCZ: 70 trials) and H (Pre-DCZ: 70 trials, Post-DCZ: 70 trials). Error bars indicate standard deviation. There was no significant difference (Wilcoxon rank-sum test; grasping time: *P* = 0.81 [Monkey A], 0.60 [Monkey H]; reaching time: *P* = 0.59 [Monkey A], 0.70 [Monkey H]). **b**, Trajectories of the index finger and thumb tips viewed from the left side during the task before and after DCZ administration; 35 days after lesioning in Monkey H. Each trajectory was plotted by the timing just before the monkey retrieved the food. Red rectangles in the middle figures indicate the areas shown in the right figures. The colour of trajectories in the right figures indicate the time from when the monkey’s thumb touched the slit. **c**, Position of the index finger and thumb tips when the monkey retrieved the food before and after DCZ administration; 35 days after lesioning in Monkey H. Line with arrows indicates the distance calculated and shown in ***d***. **d**, Distance of the thumb tip from the slit entrance at the timing when the monkeys retrieved the food before and after DCZ administration; 45 days after lesioning in Monkey A and 35 days after lesioning in Monkey H. Error bars indicate standard deviation. \**P* < 0.05 (Wilcoxon rank-sum test, Monkey A: *n* = 65 trials [Pre-DCZ], 56 trials [Post-DCZ], *P* = 0.033; Monkey H: *n* = 66 trials [Pre-DCZ], 61 trials [Post-DCZ], *P* = 1.9 × 10^-7^).

**Extended Data Fig. 3.**
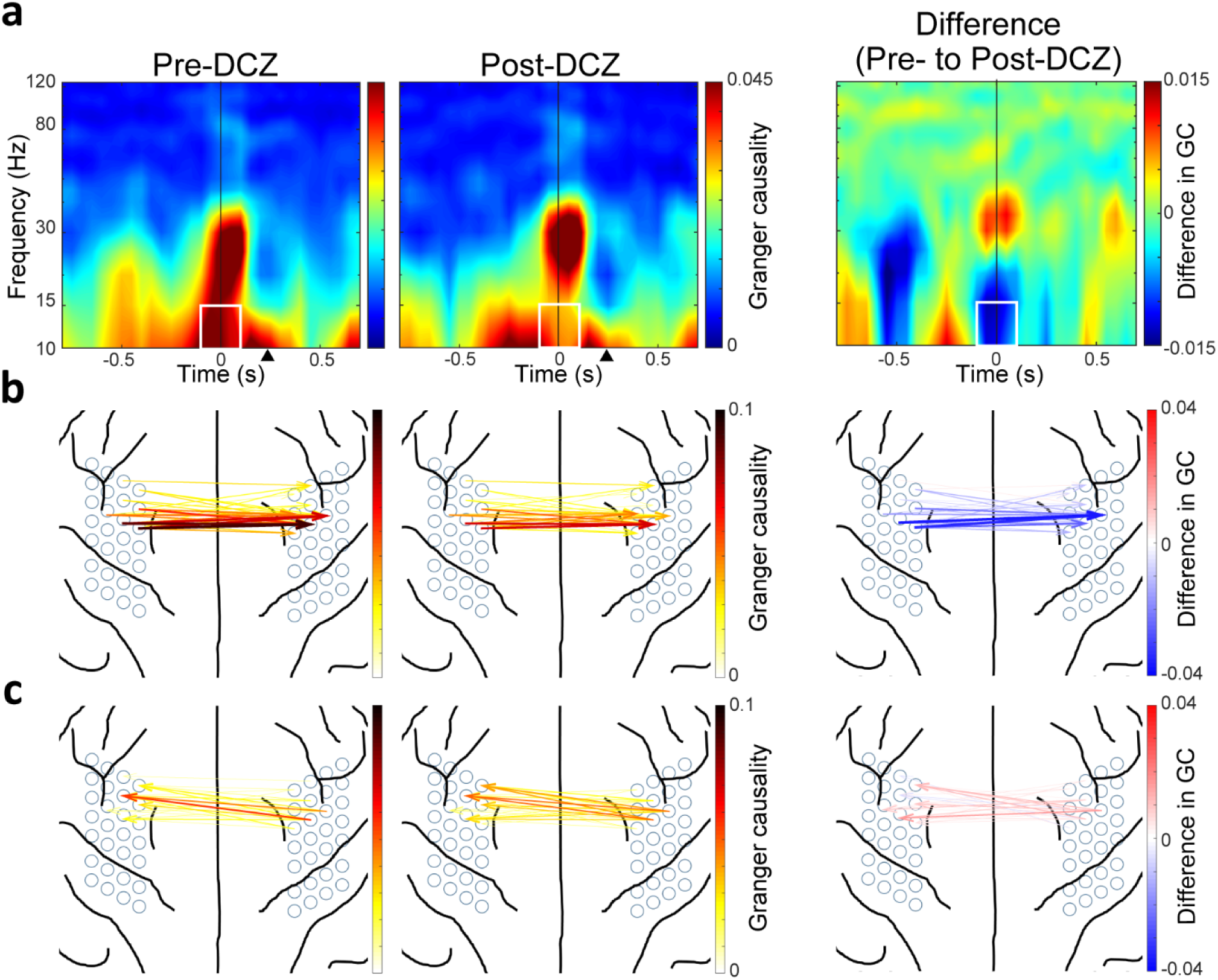
Interhemispheric connectivity is unidirectionally blocked by DREADDs (Monkey H) **a**, The averaged time-frequencygram of GC from the contralesional to ipsilesional PMd before (left figure) and after (middle figure) DCZ administration in the early recovery stage of Monkey H. Right figure shows the difference in GC between before and after DCZ administration. White rectangles indicate the time frequency ROI that was used for the calculations shown in ***b***, ***c***, and Fig. 3d. **b**, The anatomical dimension of GC from the contralesional to ipsilesional PMd in the low-frequency band around movement onset before (left) and after (middle) DCZ administration and their difference (right) (ROI shown in ***a***). **c**, The anatomical dimension of GC from the ipsilesional to contralesional PMd before (left) and after (middle) DCZ administration and their difference (right).

**Extended Data Fig. 4.**
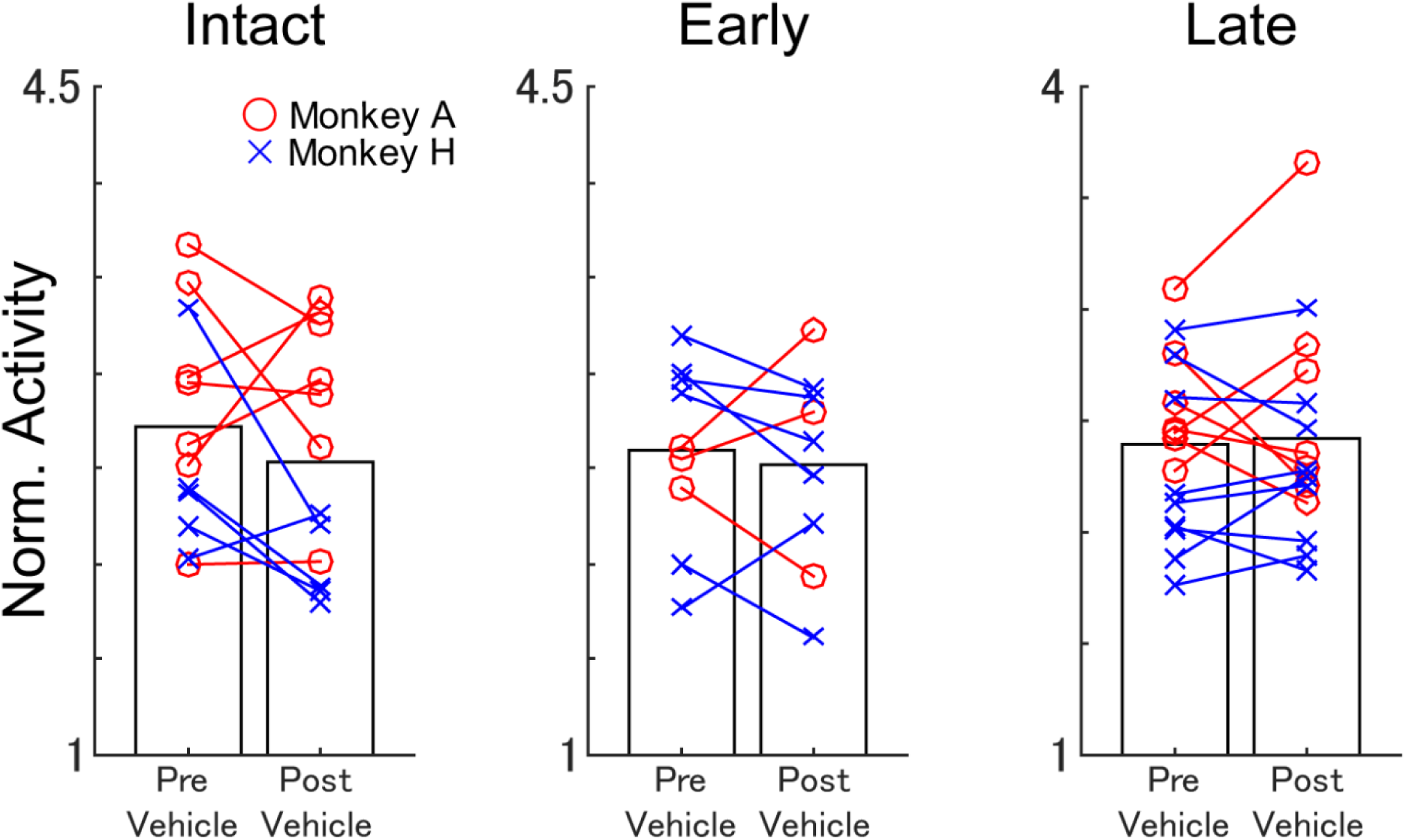
Vehicle injections do not affect the activity of the ipsilesional PM. Averaged value of normalized brain activity (ECoG) at the ipsilesional PMd before and after vehicle administration in the intact state (left), early recovery stage (middle), and late recovery stage (right) of both monkeys. There was no significant difference in activity between before and after vehicle administration in any stage. (Wilcoxon sign-rank test; intact, *n* = 12 days, *P* = 0.27; early, *n* = 9 days, *P* = 0.57; late, *n* = 16 days, *P* = 0.80.)

**Extended Data Fig. 5.**
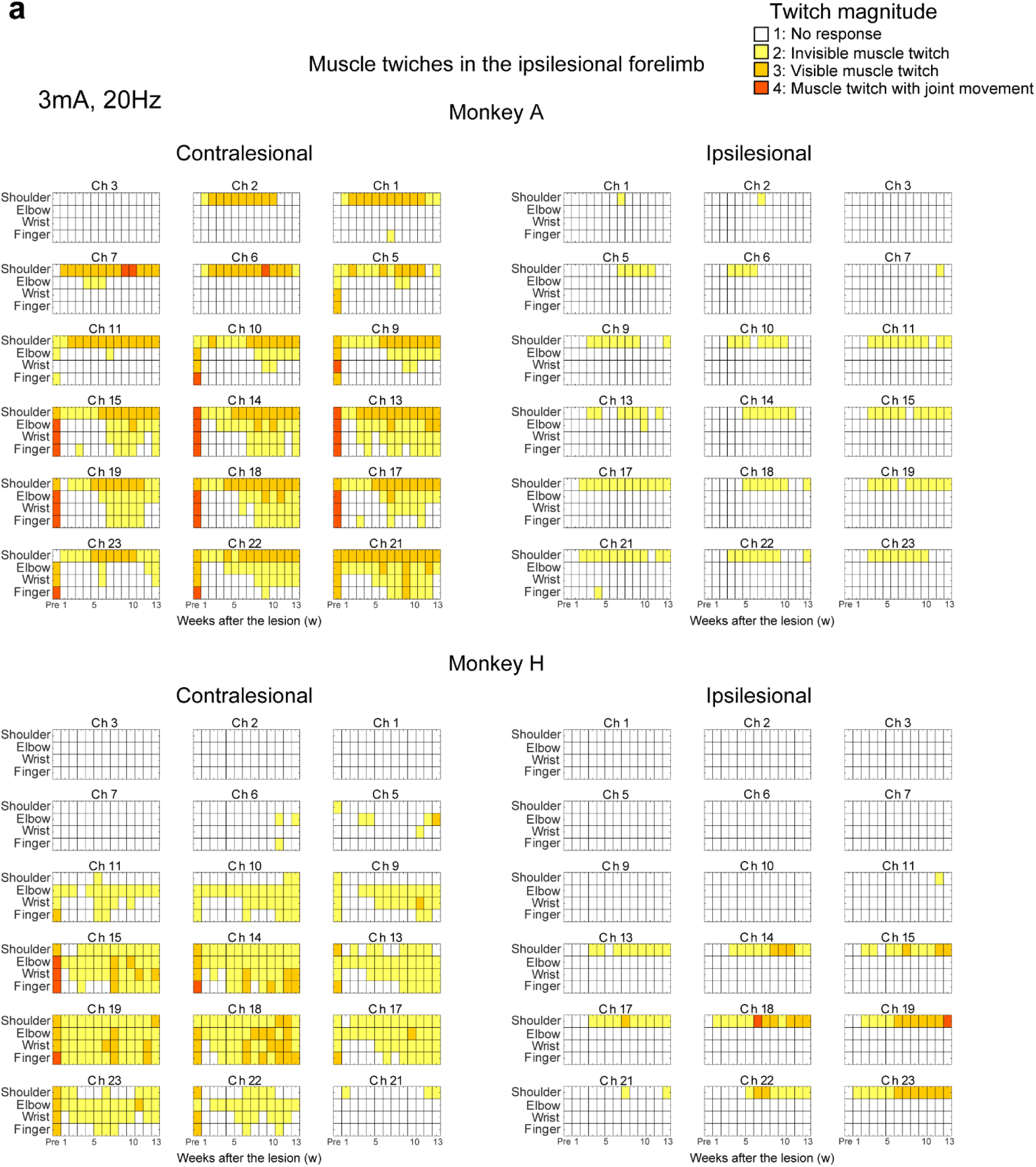
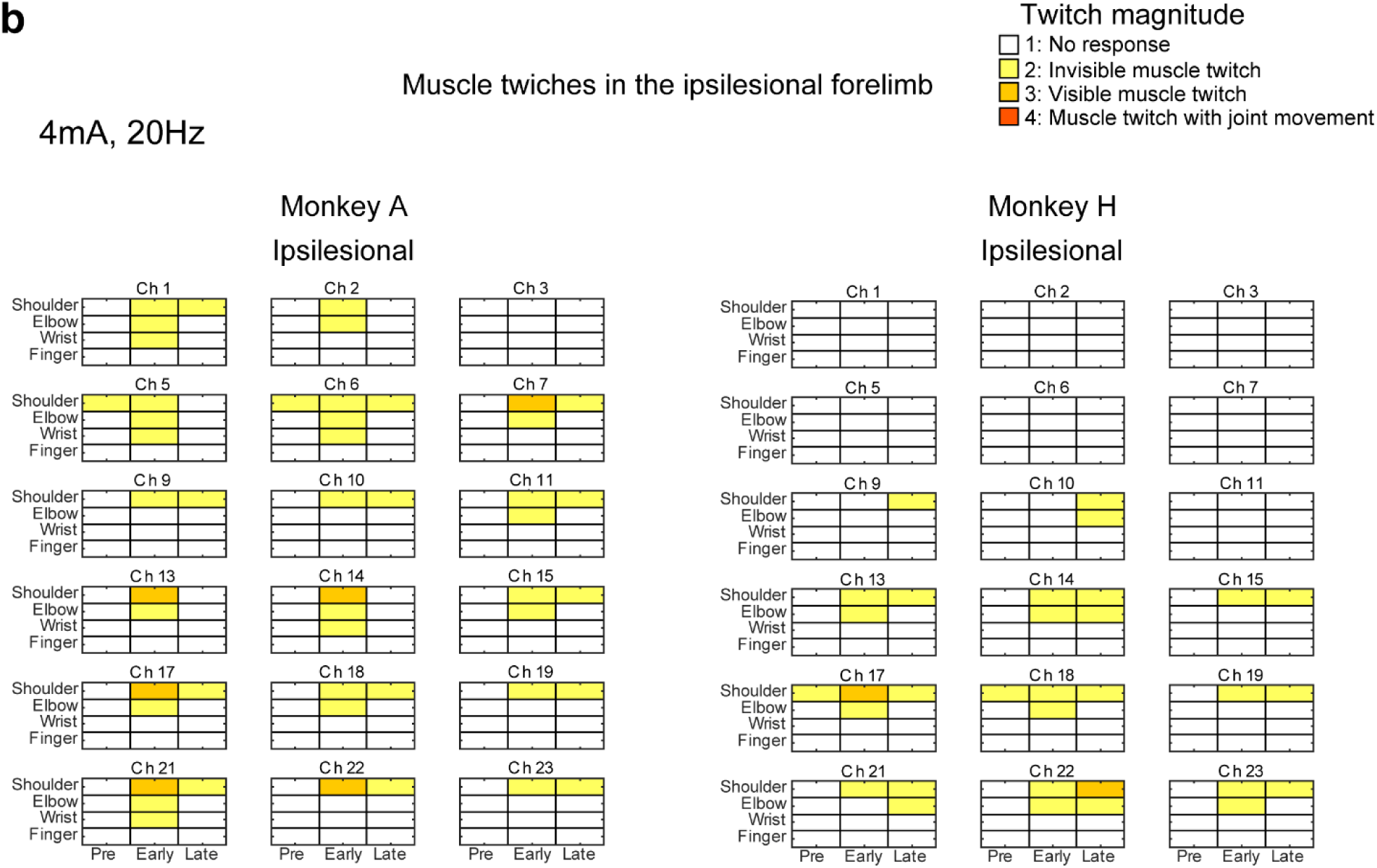
Muscle twitches induced by electrical stimulation at the ipsilesional motor cortex before and during recovery. **a**, Muscle twitches in various parts of the ipsilesional forelimb when stimulated at each channel of the ECoG electrodes on the bilateral sensorimotor cortices. Stimulation intensity was 3 mA. Colour indicates the magnitude of muscle twitches (see inset). **b**. Muscle twitches in various parts of the ipsilesional forelimb when stimulated at each channel of the ECoG electrodes on the ipsilesional cortex. Stimulation intensity was 4 mA.

**Extended Data Fig. 6.**
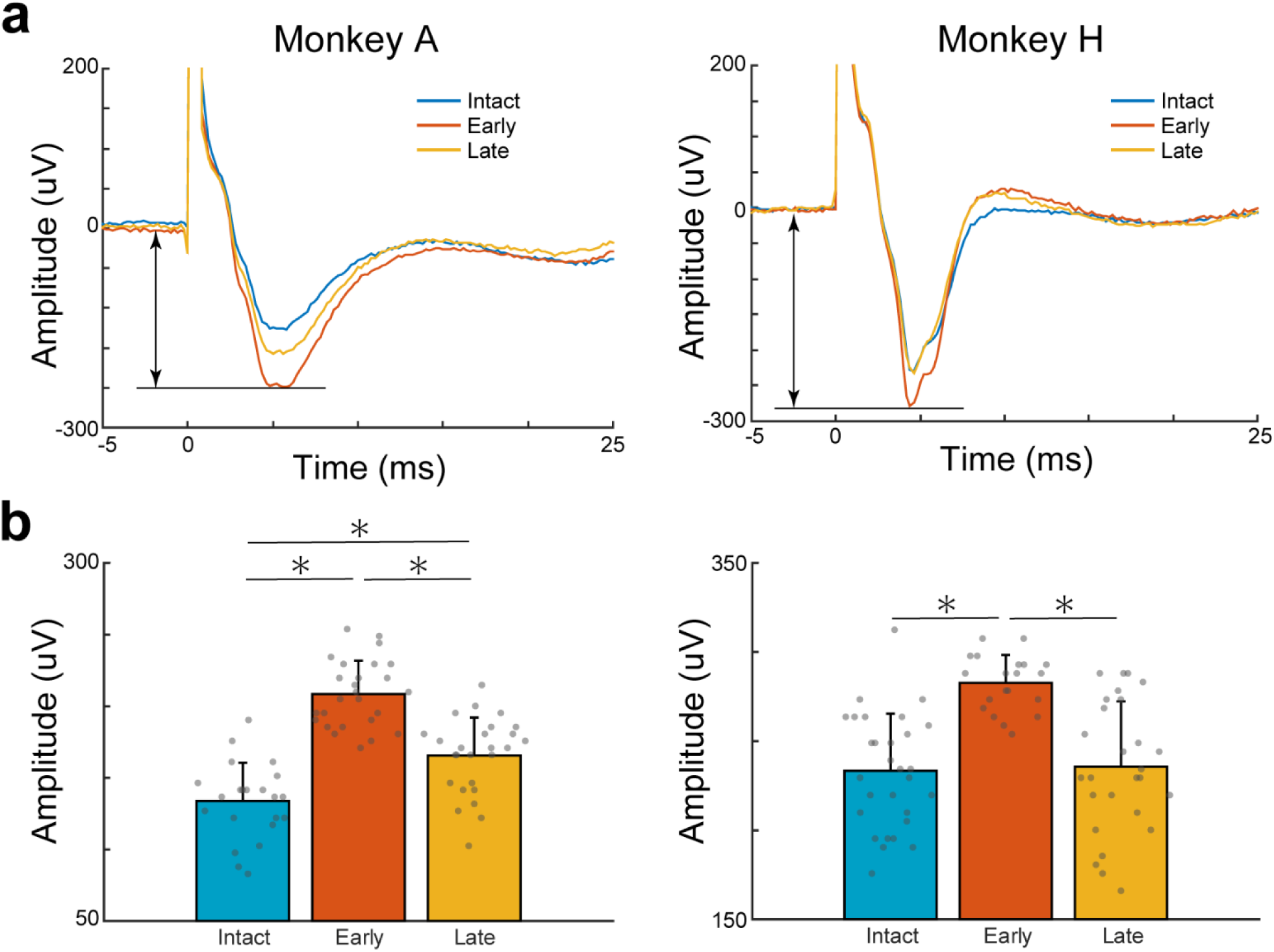
Increase in CCEPs at the ipsilesional PMd during recovery. **a**, Examples of the averaged waveform of CCEPs recorded at the ipsilesional PMd (Ch 10) when stimulated at the contralesional PMd (Ch 10) in each state (blue, intact [*n* = 25 trials, Monkey A; *n* = 29 trials, Monkey H]; orange, early [day 50, *n* = 28 trials, Monkey A; day 45, *n* = 24 trials, Monkey H]; yellow, late [day 84, *n* = 30 trials, Monkey A; day 83, *n* = 27 trials, Monkey H]). The vertical lines with arrows indicate the amplitude measured and shown in ***b***. **b**, Averaged amplitude of CCEPs recorded at the ipsilesional PMd (Ch 10) when stimulated at the contralesional PMd (Ch 10) in each state. Error bars indicate standard deviation. \**P* < 0.05 (one-way analysis of variance [*F* = 52.04, *P* < 0.05 for Monkey A; *F* = 18.33, *P* < 0.05 for Monkey H] followed by Bonferroni’s *post-hoc* test).

**Extended Data Fig. 7.**
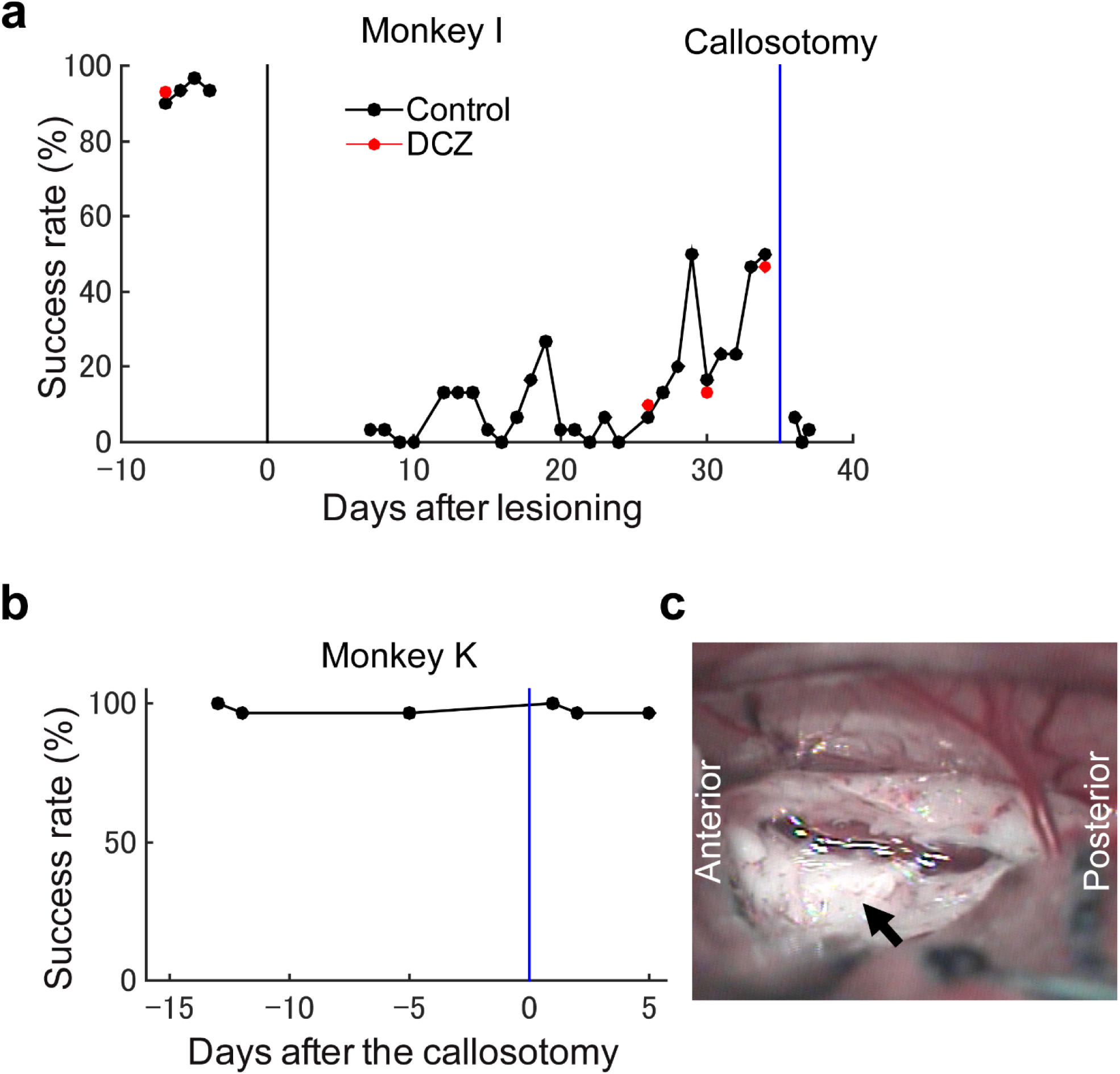
Callosotomy during recovery after l-CST lesioning impairs the recovered hand movements. **a**, Success rate of precision grip in Monkey I with partial callosotomy between the bilateral PMs during recovery from l-CST lesioning. Red circles indicate the results after DCZ administration. The black vertical line indicates the day of l-CST lesioning and the blue line indicates the day of callosotomy. **b**, Success rate of precision grip in Monkey K with partial callosotomy without l-CST lesioning. The blue line indicates the day of callosotomy. **c**, A partially dissected corpus callosum between the premotor cortex. The black arrow indicates the corpus callosum.

**Supplementary Video 1: Effect of interhemispheric pathway blockade during recovery** Video showing the recovery course of motor function in Monkey A, and the effect of interhemispheric pathway blockade with DCZ in the intact state and during recovery. In the intact state, DCZ did not affect behaviour. During recovery (Day 45), DCZ impaired the recovered finger dexterity and the monkey tried to retrieve the food repeatedly (wandering error) and dropped it frequently.

